# Human neural progenitors establish a diffusion barrier in the ER membrane during cell division

**DOI:** 10.1101/2022.02.02.478772

**Authors:** Muhammad Khadeesh bin Imtiaz, Lars N. Royall, Sebastian Jessberger

## Abstract

Asymmetric segregation of cellular components regulates the fate and behavior of somatic stem cells. Similar to dividing budding yeast and precursor cells in *C. elegans*, it has been shown that mouse neural progenitors establish a diffusion barrier in the membrane of the endoplasmic reticulum (ER), which has been associated with asymmetric partitioning of damaged proteins and cellular age. However, the existence of an ER-diffusion barrier in human cells remains unknown. Here we used fluorescence loss in photobleaching (FLIP) imaging to show that human embryonic stem cell (hESC)- and induced pluripotent stem cell (iPSC)-derived neural progenitor cells establish an ER-diffusion barrier during cell division. The human ER-diffusion barrier is regulated via Lamin-dependent mechanisms and is associated with asymmetric segregation of mono- and polyubiquitinated, damaged proteins. Further, forebrain regionalized organoids derived from hESCs were used to show the establishment of an ER-membrane diffusion barrier in more naturalistic tissues mimicking early steps of human brain development. Thus, the data provided here show that human neural progenitors establish a diffusion barrier during cell division in the membrane of the ER, which may allow for asymmetric segregation of cellular components, contributing to the fate and behavior of human neural progenitor cells.

**Summary:** Human neural progenitors (NPCs) establish a diffusion barrier during cell division in the membrane of the endoplasmic reticulum, allowing for asymmetric segregation of cellular components, which may contribute to the fate and behavior of human NPCs.

## Introduction

Somatic stem cells can self-renew via symmetric, duplicating division or divide asymmetrically to produce daughter cells of different cell fate. Asymmetric cell division of stem cells is associated with asymmetric segregation of cellular components (Moore and Jessberger, 2017). Unequal partitioning of cell content was for example shown in stem cells of *D. melanogaster* where the cell fate determinant NUMB becomes asymmetrically inherited, hence generating daughter cells with distinct fates (Cayouette and Raff, 2002; Rhyu et al., 1994). Asymmetric segregation also extends to other cell cargoes such as mitochondria, damaged proteins, and centrioles (Katajisto et al., 2015; McFaline-Figueroa et al., 2011; Wang et al., 2009). Asymmetric inheritance of cargoes may mediate advantages to the stem cell, whose health must be maintained to conserve its proliferative ability (Henderson and Gottschling, 2008). Further, asymmetrical segregation of certain cargoes is essential for the activation and function of daughter cells. For example, lysosome inheritance has been shown to correlate with the activation of hematopoetic stem cells (Loeffler et al., 2019). However, under certain conditions associated with cancer but also aging, proper asymmetric segregation of cargoes appears to become impaired, resulting in the disruption of tissue homeostasis and maintenance (Knoblich, 2010).

The mechanisms underlying asymmetric segregation have been studied in a variety of organisms from budding yeast, *D. melanogaster, C. elegans* to mice (Lee et al., 2016; Luedeke et al., 2005; Moore et al., 2015). Contributing to asymmetric cell division, a diffusion barrier in the membrane of the endoplasmic reticulum (ER) has been identified and proposed as a critical mediator determining the segregation of aging factors (Clay et al., 2014; Shcheprova et al., 2008). In mouse neural progenitor cells, the ER membrane diffusion barrier becomes weakened with age, resulting in increased symmetric inheritance of damaged proteins with advancing age (bin Imtiaz et al., 2021; Moore et al., 2015). Even though a barrier in the ER membrane has been described in a variety of species, its presence and relevance for human neural progenitor cells (NPCs) remains unknown.

Using human embryonic stem cell (hESC)- and induced pluripotent stem cell (iPSC)-derived progenitor cells, we show that human NPCs establish a diffusion barrier in the ER membrane during cell division that is regulated via Lamin-dependent mechanisms and that is associated with the asymmetric segregation of damaged proteins. Further, we used genome-edited, hESC-derived forebrain organoids to show the presence of an ER-membrane diffusion barrier during more naturalistic human NPC divisions.

## Results & Discussion

### ER membrane diffusion barrier in human progenitors

To study whether a diffusion barrier in the ER membrane is present in human NPCs, we used fluorescence loss in photobleaching (FLIP) experiments, in analogy to experiments previously described in mouse NPCs (Figure 1A) (bin Imtiaz et al., 2021; Moore et al., 2015). First, we developed tools to visualize the human ER membrane and lumen in human NPCs cells. Membrane markers used previously were toxic to human NPCs and resulted in high rates of cell death. Hence, we tested a plethora of proteins localized to the ER membrane and eventually used Suppressor/Enhancer of Lin-12-like (SEL1L) tagged with green fluorescent protein (GFP) as a novel marker for the ER membrane (bin Imtiaz et al., 2021). Human progenitors derived from hESCs were transfected with fluorescent proteins targeted to the lumen (KDEL-GFP, hereafter called LumER-GFP) or to the membrane of the ER (Sel1L-GFP, hererafter called MemER-GFP).

**Figure 1.**
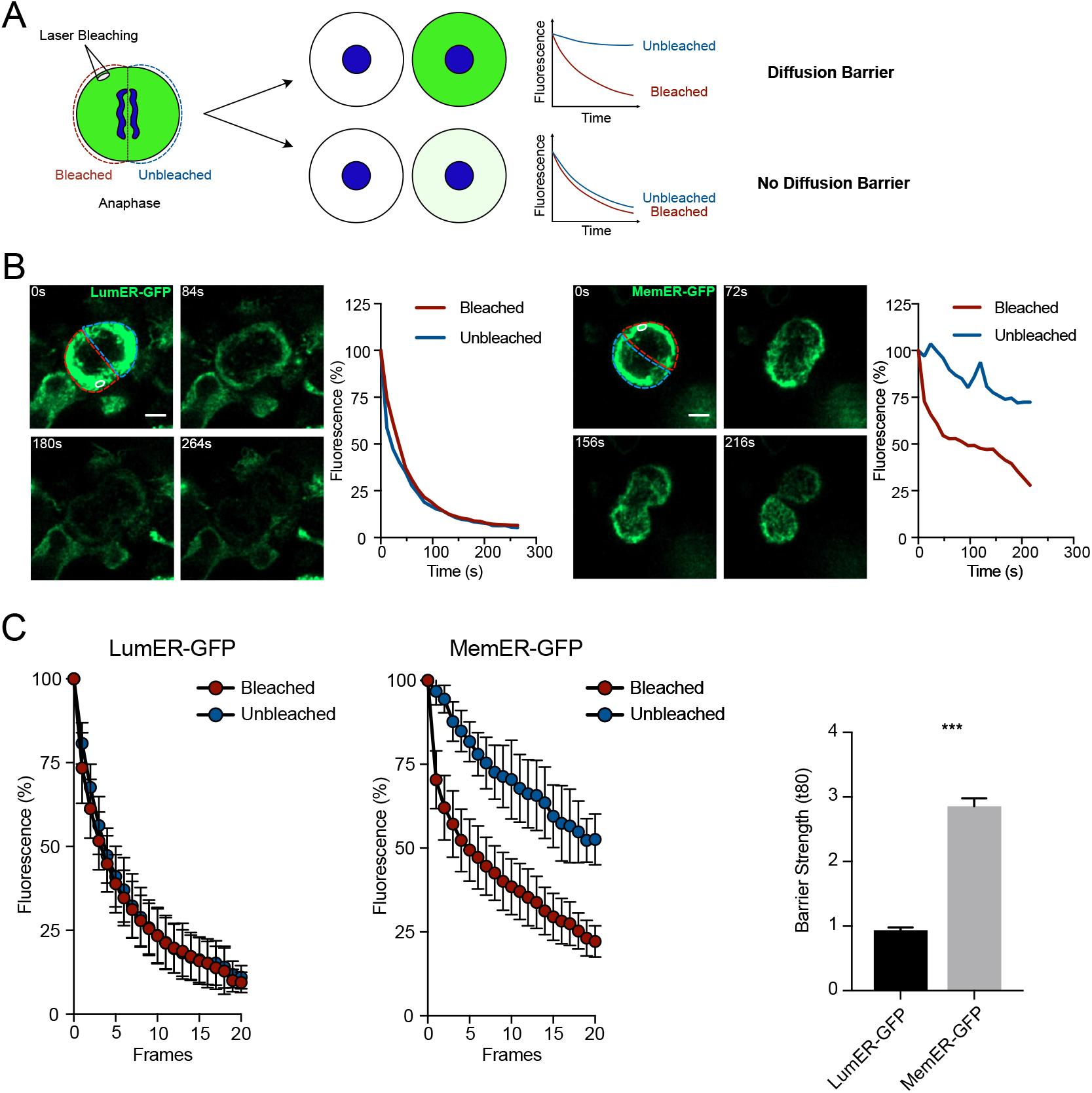
An ER membrane diffusion barrier in human neural stem cells. (A) An outline of the FLIP assay. A region is selected and continuously bleached in cells expressing a marker for a GFP marker. The compartment where the bleached region is located (red outline) is termed “Bleached” whereas the unbleached compartment (blue outline) is termed “Unbleached”. The region is continuously bleached until no visible GFP signal is present between the two compartments. The fluorescence is then measured in both Bleached and Unbleached during the course of bleaching. In presence of a diffusion barrier, loss of fluorescence is only limited to Bleached whereas in the absence of diffusion barrier (above), loss of fluorescence is equal in both Bleached and Unbleached (below). (B) Single cells expressing either Lum-GFP (right) or MemER-GFP (left) undergoing FLIP assays are shown. The region bleached is highlighted (white). The Bleached and the Unbleached compartments of the cells are shown (red and blue respectively). Measured fluorescence intensities for the two compartments are plotted against time (right). (C) Average fluorescence intensities for the Bleached and the Unbleached regions are shown for LumER-GFP and MemER-GFP (left). Barrier strength indices of both Mem-GFP and LumER-GFP are also plotted (right). Scale bars represent 5μm (B). ***p < 0.001.

To probe for the existence of a diffusion barrier in human progenitor cells, we used FLIP experiments during cell division (Figure 1A). Once cells entered anaphase, a region was selected in one compartment of the dividing cell and bleached continuously. While cells progressed through cell division, the fluorescence signal in the two compartments was measured and plotted with the intensities normalized to the pre-bleach signal in both compartments. In the event a diffusion barrier limiting free movement of the tagged protein between the two compartments, the unbleached compartment will lose significantly less fluorescence signal. In the absence of a diffusion barrier, the decay in the intensity of the signal in both compartments will be equal (Figure 1A). Fluorescence intensity of LumER-GFP showed no difference in the loss of GFP signal between the bleached and unbleached compartments, indicating that the ER is continuous, and that diffusion is not limited within the lumen of the ER (Figure 1B-C, Movie S1). In contrast, we found a localized loss of fluorescence in the bleached compartment while the unbleached compartment retained most of its GFP signal when we performed FLIP experiments using MemER-GFP (Figure 1B-C, Movie S1). These results show the presence of a diffusion barrier within the membrane of the ER during division, limiting free diffusion of proteins in the continuous ER membrane of human NPCs (Figure 1B-C). To corroborate the findings of a hESC-derived NPC line, we next tested a human iPSC-derived NPC line for the presence of an ER membrane diffusion barrier (Bohaciakova et al., 2019). Again, we found that iPSC-derived NPCs established a diffusion barrier in the ER membrane during anaphase (Figure S1A-B).

These results show that human NPCs establish a diffusion barrier that restricts free diffusion of ER membrane proteins during mitosis. Similar mitotic ER membrane diffusion barriers have been reported in yeast, mice and *C. elegans*, and our results suggest that this is a conserved mechanism between stem cells of different species (Lee et al., 2016; Luedeke et al., 2005; Moore et al., 2015).

### Lamin-dependent regulation of human ER barrier strength

Next, we aimed to modulate the strength of the human ER membrane diffusion barrier. Lamins, which are intermediate filaments in the nucleus, have been implicated in the weakening of the diffusion barrier that occurs with advancing age (bin Imtiaz et al., 2021). Progerin, which is a mutant form of lamin A, implicated in Hutchinson-Gilford progeroid syndrome (HGPS), has been previously used to mimic certain cellular aging phenotypes in vitro (Miller et al., 2013; Moore et al., 2015). To determine whether the human ER membrane diffusion barrier is regulated with progerin-induced aging, we overexpressed progerin in human NPCs (Figure 2A). We then performed FLIP assays on MemER-GFP hNPCs and observed that progerin expression weakened the strength of the ER membrane diffusion barrier in human NPCs (Figure 2B-C, Movie S2), suggesting that nuclear lamins play a role in the establishment and regulation of the ER membrane diffusion barrier in human NPCs.

**Figure 2.**
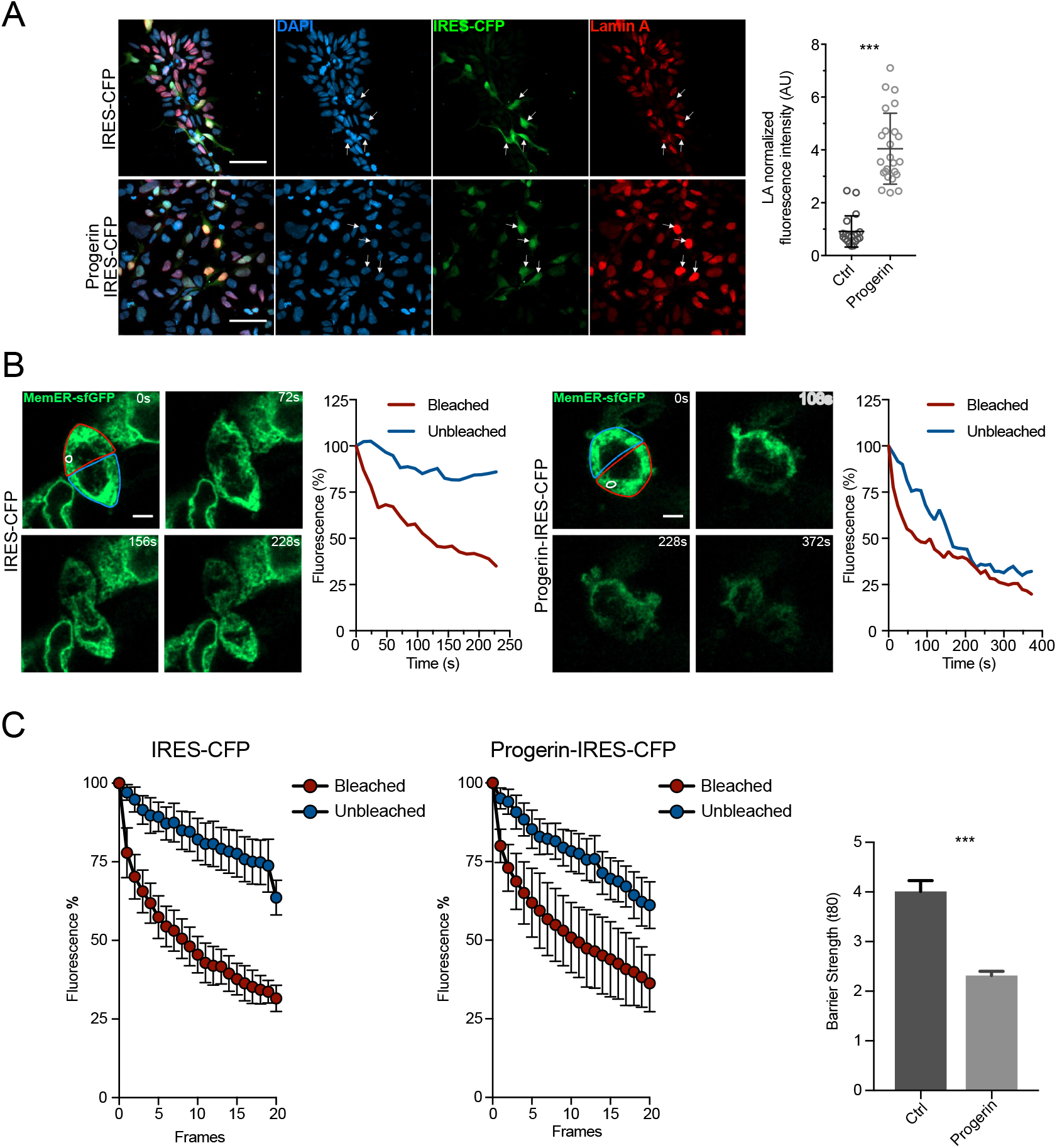
Progerin expression weakens the ER membrane diffusion barrier. (A) Representative images showing the expression of progerin in hNPCs. hNPCs were transduced with either a control (IRES-CFP) or progerin (Progerin IRES-CFP) plasmid and were then stained against DAPI (blue), IRES-CFP (green) and lamin-A (red). (B) Single cells expressing MemER-GFP and either control (IRES-CFP; right) or progerin (Progerin-IRES-CFP; left) undergoing FLIP assays are shown. The region bleached is highlighted (white). The Bleached and the Unbleached compartments of the cells are shown (red and blue respectively). Measured fluorescence intensities for the two compartments are plotted against time (right). (C) Average fluorescence intensities for cells expressing either control (IRES-CFP) or progerin (Progerin-IRES-CFP) that underwent FLIP assays are shown. Bleached (red) and Unbleached (blue) fluorescence intensity averages are shown over time (s; left). Barrier strength indices of control and progerin cells are also plotted (right). Scale bars represents 50μm (A) and 5μm (B). ***p < 0.001.

### Asymmetric segregation of damaged proteins

Previous studies suggested a correlation between the strength of the ER membrane diffusion barrier and asymmetric segregation of mono- and poly-ubiquitinated proteins, i.e., damaged proteins (Clay et al., 2014; Moore et al., 2015). Thus, we next tested whether ubiquitinated proteins are asymmetrically segregated during human NPC divisions. Using immunostaining to label mono- and poly-ubiquitinated proteins, we found that human NPCs showed asymmetric segregation of ubiquitinated proteins (Figure 3A). We next analyzed if a weakened ER membrane diffusion barrier, induced by overexpression of progerin, affects asymmetric segregation of ubiquitinated proteins. Indeed, progerin expressing cells, associated with a weakened ER membrane diffusion barrier, showed increased symmetric segregation of ubiquitinated proteins compared to control cells (Figure 3B), suggesting that the ER membrane diffusion barrier is involved in the asymmetric segregation of ubiquitinated proteins in human NPCs.

**Figure 3.**
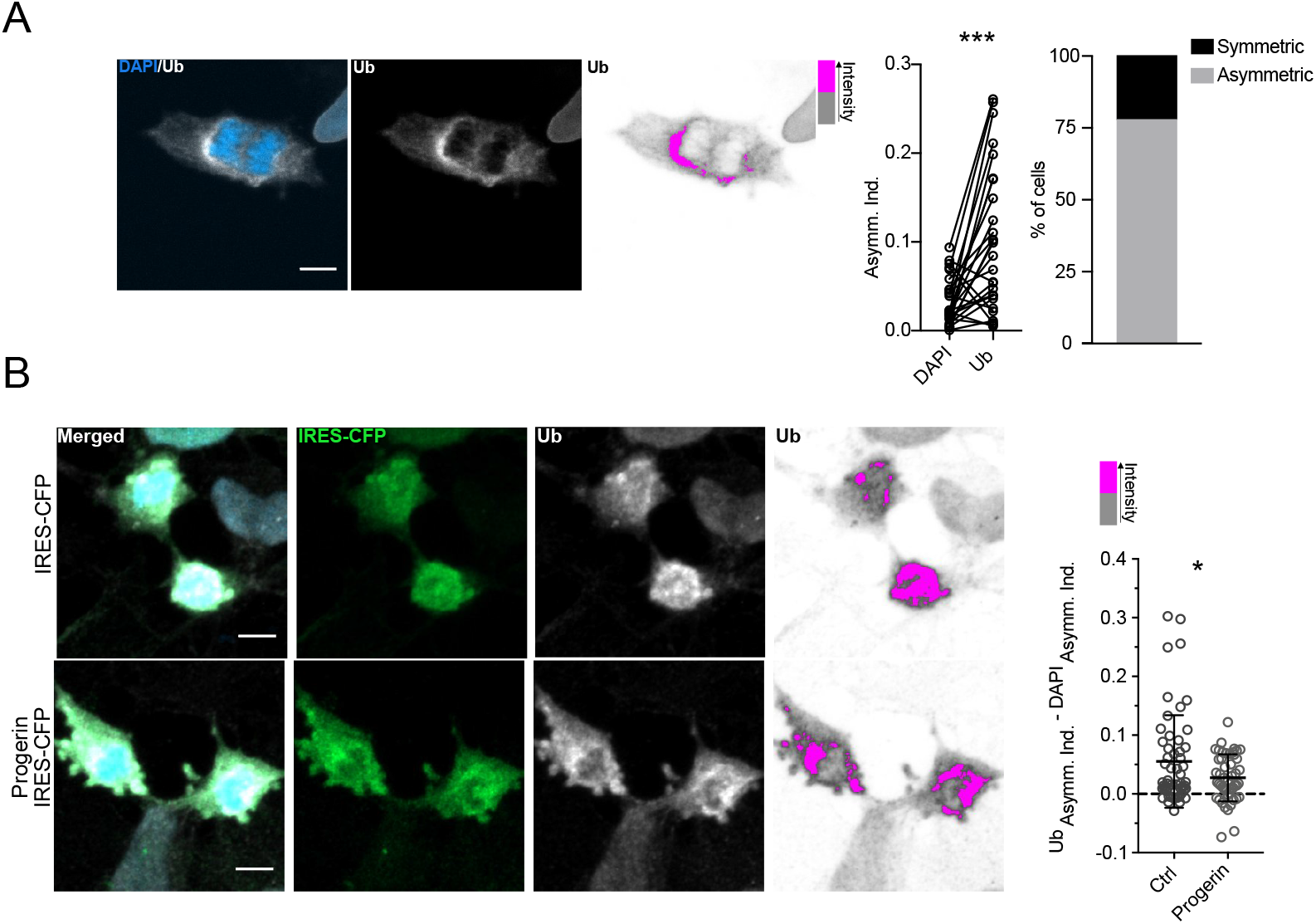
Ubiquitinated proteins are asymmetrically segregated in hNPCs. (A) A representative image showing asymmetrical segregation of ubiquitinated proteins. The hNPCs were stained for DAPI (blue) and ubiquitin (white; left). High ubiquitin intensity in this example is shown in pink whereas low ubiquitin intensity is shown in grey. Asymmetry indices of DAPI and ubiquitin were then calculated for each cell and plotted (middle). Cells with a higher asymmetry index of ubiquitin were classified as asymmetric and plotted (right). (B) Representative images of hNPCs expressing either control (IRES-CFP) or progerin (Progerin-IRES-CFP) showing asymmetric segregation of ubiquitinated proteins. hNPCs were stained for DAPI (blue), IRES-CFP (green) and ubiquitin (white). High ubiquitin intensity is shown in pink whereas low ubiquitin intensity is shown in grey. The difference between asymmetry indices of Ub and asymmetry indices of DAPI is plotted (right). Scale bars represent 5μm (A and B). *p < 0.05, ***p < 0.001.

### ER membrane diffusion barrier in neural progenitors of regionalized forebrain organoids

Following our *in vitro* studies on 2-dimensional cultured cells, we next aimed to study whether human NPCs establish a diffusion barrier in more naturalistic, 3-dimensional *in vitro* models of the developing human brain, i.e., regionalized forebrain organoids. To obtain forebrain organoids allowing for FLIP-based experiments, we first established hESC lines that constitutively expressed either MemER-GFP or LumER-GFP from the human safe harbor site AAVS1 (Chen et al., 2015). We mixed hESCs positive for MemER-GFP or LumER-GFP with non-labelled hESCs, to produce mosaic human forebrain organoids that enabled FLIP to be performed on individual cells of the densely packed ventricular zone, consisting largely of SOX2-expression NPCs that orient themselves around ventricle-like structures within cortical units (Figure 4A; Figure S2A; Denoth-Lippuner et al., 2021; Qian et al., 2016). We observed a mosaic distribution of GFP positive cells throughout the organoids, indicating that the constitutive expression of the tagged markers did not confer a disadvantage to labeled cells. We directly observed cell division events of NPCs in organoids using time-lapse imaging over 4 hours (Figure 4A-B).

**Figure 4.**
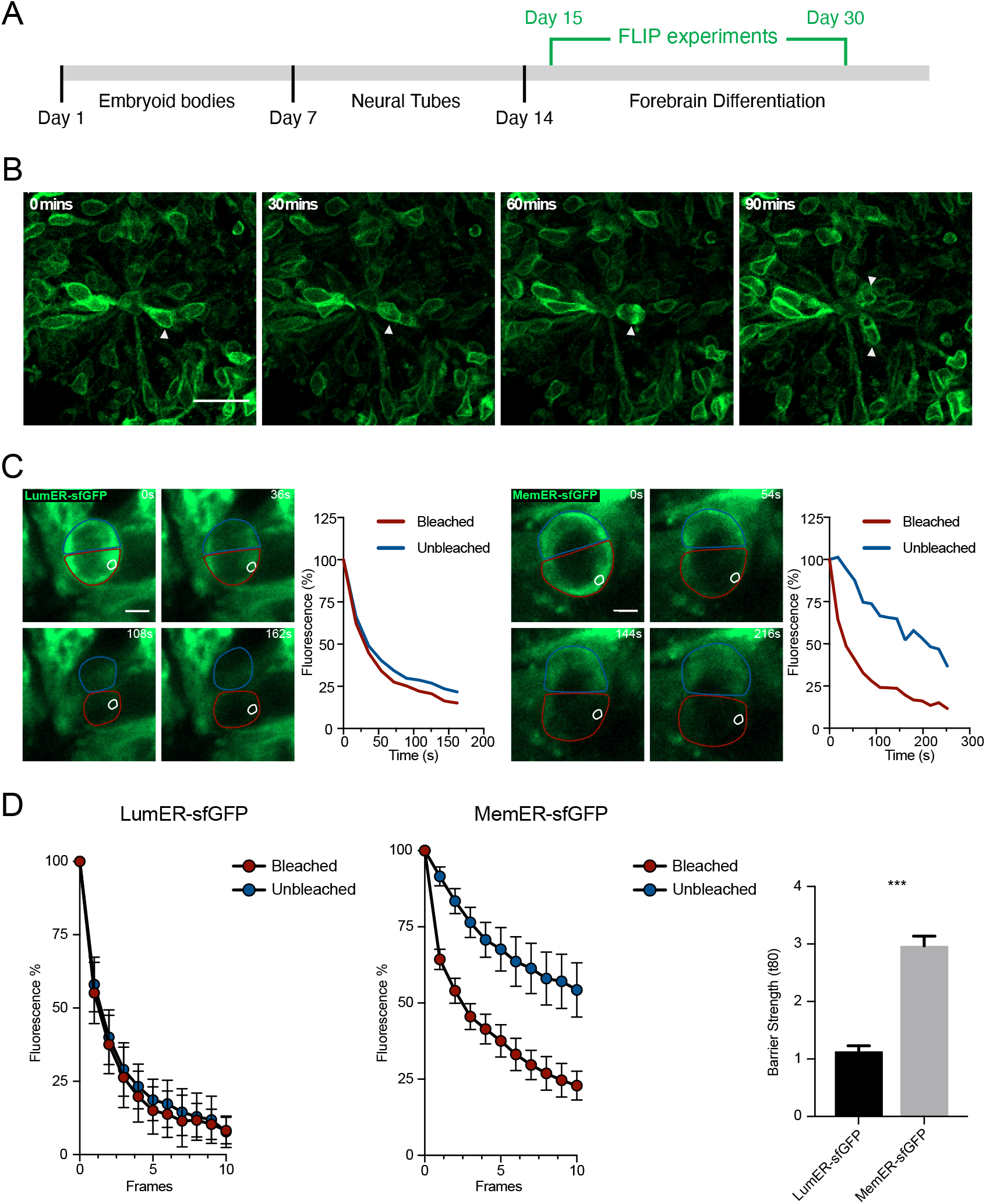
An ER membrane diffusion barrier is established in dividing neural progenitors of human forebrain organoids. (A) The experimental outline of the FLIP assays on forebrain organoids is shown. Embryoid bodies from human embryonic stem cells (hESCs), which stably expressed either the LumER-GFP or MemER-GFP, were made. This was followed by induction of neural tubes and then organoids mimicking human forebrain differentiation. FLIP assays were performed between 15 and 30 days post formation of embryoid bodies. (B) A neural progenitor division in a MemER-GFP organoid is shown. The neural progenitor (arrowhead) performs interkinetic nuclear migration towards the ventricular unit before undergoing mitosis. The two daughter cells are also shown (arrowheads). (C) Single cell traces of LumER-GFP and MemER-GFP progenitors undergoing mitosis in organoids. An area (white) was selected and bleached continuously from the onset of anaphase until no GFP signal was visible between the two compartments. The fluorescence intensities of the bleached (red) and unbleached (blue) are plotted (right) for each cell. (D) Average fluorescence intensities of the two compartments for both LumER-GFP and MemER-GFP cells are plotted against frames (left). Barrier strength indices for LumER-GFP and MemER-GFP is also plotted (right). Scale bars represent 20μm (B) and 5μm (C). ***p <0.001.

We then performed FLIP assays of dividing NPCs in LumER-GFP and - MemER-GFP-labeled organoids (Figure 4C, Movie S3). In analogy to 2D cultured NPCs, we found that LumER-GFP cells displayed equal loss of fluorescence in the two compartments during division, indicating that the ER membrane is continuous and that LumER-GFP diffuses freely between the two compartments (Figure 4C-D). However, cells expressing MemER-GFP displayed reduced fluorescence loss in the unbleached compared to the bleached compartment, clearly indicating the establishment of an ER membrane diffusion within dividing NPCs in hESC-derived organoids (Figure 4C-D). When the unbleached and bleached fluorescence intensities were subtracted for individual cells, we observed an ER diffusion barrier in the majority of cells expressing MemER-GFP (Figure S2B-C). Thus, our results identify a diffusion barrier in the ER membrane as a mechanism for establishing asymmetry in human NPCs during cell divisions.

These results shown here demonstrate the presence of an ER membrane diffusion during human NPC divisions. Similar diffusion barriers in the membrane of the ER have been reported previously in yeast, mouse and *C. elegans* cells (Lee et al., 2016; Luedeke et al., 2005; Moore et al., 2015; Saarikangas et al., 2017). Our results suggest that the establishment of an ER membrane diffusion barrier represents a conserved mechanism between stem cells of different species. A weakening of the ER barrier with progerin overexpression indicates that lamin-dependent mechanisms are important regulators of barrier strength, as previously described in mouse cells (bin Imtiaz et al., 2021; Moore et al., 2015). However, the molecular identity of the ER barrier in mammalian cells remains poorly understood. This is in contrast to budding yeast where substantial progress has been made over the last years to characterize the molecular composition and functional properties of diffusion barriers in the ER membrane during cell divisions (Clay et al., 2014; Megyeri et al., 2019). In human NPCs, the specific cargoes that the barrier in the ER membrane partitions remain largely unknown even though our data found a correlation between asymmetric segregation of mono- and polyubiquitinated proteins with barrier strength. Future studies will need to identify if disruption or weakening of the ER barrier affects human NPC behavior and early steps of human brain development. The data shown here represent the foundation for future experiments with the aim to further our understanding of the role of asymmetric segregation during human progenitor cell divisions.

## Materials & Methods

### 2D cell culture and forebrain organoids

Both lines of human NPCs were plated on Poly-Ornithine (Sigma Aldrich) and Laminin (Sigma Aldrich) coated plates. The media for human NPCs obtained from Roche was supplemented with B-27-supplement minus vitamin A (Thermo Fisher), N2-supplement (Thermo Fisher), Neurobasal (Thermo Fisher), EGF-2 (PeproTech), FGF (PeproTech) and DMEM/F-12 GlutaMAX (Thermo Fisher). The human NPCs sourced from Polymenidou group (UZH) were plated in media supplemented with DMEM/F12 (Thermo Fisher), 0.5X B27-supplement (Thermo Fisher), 0.5X N2 supplement (Thermo Fisher), 1X GlutaMAX (Thermo Fisher) and 20 ng/ml bFGF Gibco #PHG0261. Both lines were passaged with 0.05% Trypsin (Thermo Fisher) that was then blocked with an equal volume of Defined Trypsin Inhibitor (Thermo Fisher). The media was changed every 2 days. Human NPCs were counted and 4×10^6^ were used for electroporations. Cells were then resuspended in 100μl nucleofactor solution (Lonza) that contained 3μg of the plasmids. The cells were then electroporated using the AMAXA electroporation system (Lonza) using the A-033 program. The cells were then plated in poly-Ornithin (Thermo Fisher) and Laminin (Thermo Fisher) coated plates. H9 hESC were maintained in feeder-free conditions without antibiotics in mTeSR™ Plus (Stem Cell Technologies 05825) on hESC qualified Matrigel (Corning, 354277) coated plates. Routine passaging was performed with ReLeSR™ (Stem Cell Technologies, 05872). 10μM Y-27632 (Stem Cell Technologies, 72302) added to media post passaging. For long term storage, hESCs were stored at −170°C in CyroStore® CS10 (Sigma, C2874). All experiments with hESCs received approval from the Canton of Zurich’s Kantonale Ethik-Kommision (KEK).

Electroporation was conducted with an Amaxa nucleofector II using program A-23 according to the manufacturer’s guidelines. Prior to electroporation, hESCs were treated with mTeSR™ Plus containing 10μM Y-27632 for at least 2 hours. 2 million cells were passaged with Accutase (Sigma, A6964) and were resuspended in Nucleofector V (Lonza, VCA-1003). Positive cells were selected for either Neomycin resistance using 100μg/ml G418 sulfate (Gibco, 10131035). Organoids were produced with a mix of WT and GFP expressing hESC at a ratio of 1:1 in order to reduce background fluorescence. Human forebrain organoids were generated via an adapted protocol (Qian et al., 2018). Alternations are as follows: hESCs were grown in feeder free conditions and at day 0 were passaged to get single cells. Single cells were aggregated in AggreWell™800 (Stem Cell Technologies, 34811) following the manufactures instructions. Embryoid bodies were harvested the following day and maintained in mTeSR™-E5 (Stem Cell Technologies, 05916) with 2μM Dorsomorphin (Sigma, P5499) and 2μM A83-01 (Tocris, 2939). From day 4 we followed the protocol described in Qian et al. 2018. Organoids were fixed for 30 minutes in 4% PFA and were stored in 30% sucrose until further use.

### Cloning and constructs

Sel1L sequence (corresponding to amino acids 178-310 from AAH57452.1; gift from M. Molinari) and KDEL was cloned into a retroviral, GFP expressing vector (Addgene 16664). Progerin construct (Addgene 17663) was cloned into a retroviral backbone vector that contained a CAG promoter followed by an internal ribosome entry site (IRES) and CFP. For control plasmid, the same plasmid without progerin cloned in was used. Flpe-ERT2 was removed from AAVS1-Neo-CAG-Flpe-ERT2 (Addgene 68460) via restriction digest and replaced with Se1L-GFP or KDEL-GFP.

### Retrovirus transduction

90,000 human NPCs were plated in 12-well plates coated with Poly-L-Ornithine (Sigma Aldrich) and Laminin (Sigma Aldrich) and incubated at 37°C overnight. Next day, the media was changed in the morning. Six hours after the media change, 500μl of the media was transferred to a tube and 1μl of the corresponding virus was added.

### Image acquisition

Zeiss LSM800 were used to acquire the images. ImageJ/Fiji was used to quantify all the images. The images were blinded before the counting/analysis was performed.

### Fluorescence Loss in Photobleaching (FLIP) assays

90,000 human NPCs were plated in a well of Chambered Coverglass wells that had been coated with Poly-L-Ornithine (Sigma Aldrich) and Laminin (Sigma Aldrich). 63x, oil-immersive objective was used to do the following imaging. For hNPCs, the wells were manually scanned to identify GFP-positive cells in mitosis. Upon identification, the cells were marked and once they started undergoing anaphase, a region of interest was selected close to the future cleavage furrow. A prebleach image was taken before a bleach with 80 iterations and 3% laser power was applied. This bleaching was applied over 8 seconds and the next image was then acquired. There was a 12 second interval between each subsequent image, with 8 seconds being used for the bleach. The cell was continuously bleached in this manner until no GFP signal was visible between the two compartments of the dividing cell. Then, the fluorescence intensities in the bleached and the unbleached compartments were measured. The fluorescent intensities for both compartments was then divided by the area. The background intensities were subtracted from both bleached and unbleached intensities. All the measured intensities were normalised to the pre-bleach image that had been set to 100%. For the average intensity graphs, fluorescent intensities in both compartments was averaged across all cells (Figure 1C, 3C and S1B).

Organoids were embedded into Phenol Red-free, growth factor reduced Matrigel (Corning) on chambered coverglasses that was allowed to polymerise for 30 minutes at 37°C. Imaging media was then added prior to performing FLIP assays. For FLIP on the organoids, a 20X objective with a digital zoom of 2x was used. The positions of cortical units were marked and scanned manually in the GFP channel to identify cells undergoing mitosis. Once anaphase commenced in the identified cells, a prebleach picture was acquired. This was followed by selecting and bleaching a region close to the future cleavage furrow. The bleach was performed with 80 iterations and 3% laserpower. The time interval between images was set at 18 seconds where 12 seconds were needed to perform the required bleaching. The cells were continuously bleached until there was no GFP signal visible between the two compartments of the dividing cell. Then, the fluorescence intensities in the bleached and the unbleached compartments were measured. The fluorescent intensities for both compartments was then divided by the area. The background intensities were subtracted from both bleached and unbleached intensities. All the measured intensities were normalised to the pre-bleach image that had been set to 100%. For the average intensity graphs, fluorescent intensities in both compartments was averaged across all cells (Figure 4D).

The barrier strengths were calculated by first plotting the fluorescence intensities from both the bleached and unbleached compartments in PRISM. These were then each fitted to a one-phase decay with the following constraints: Plateau = 0, Y0 = 100, and K > 0. The value of the constant K was determined from the fit in Prism and substituted into the following equation to calculate the time it takes for the corresponding compartment to fall to 80% from the initial 100% of fluorescence signal.

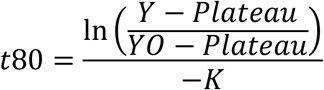

In this, Y = 80, Y0 = 100, Plateau = 0 and K was derived from the corresponding fit in Prism.

To calculate the barrier strengths, the t80 for unbleached was divided by t80 for bleached.

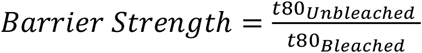

To calculate the standard error in each compartment, the following equation was used.

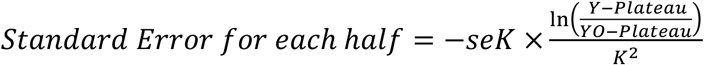

Where seK is the standard error of the constant K, derived from Prism.

For the total standard error, the following equations were used.

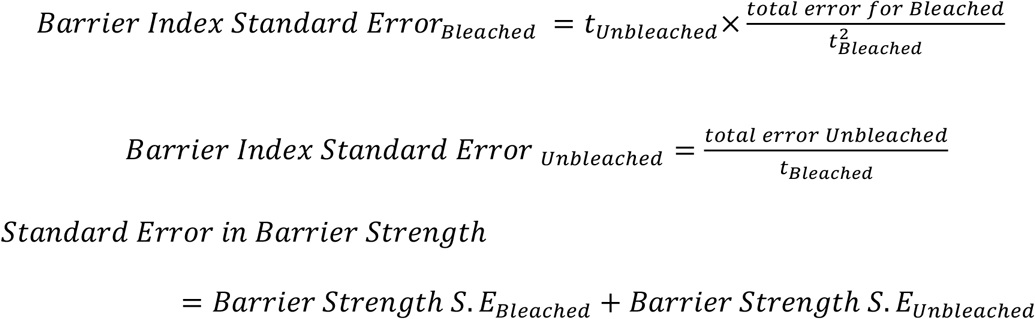

This analysis has been previously described (bin Imtiaz et al., 2021).

### Immunostaining

For immunostaining, coverslips were first coated with Poly-L-ornithine (Sigma Aldrich) for 1 hour at 37°C and then overnight with Laminin (Sigma Aldrich) at 4°C. 90,000 human NPCs were plated on the coated coverslips. They were then fixed with 4% PFA (Sigma Aldrich) for 15 mins at room temperature before being washed 3 times with PBS. After fixation, the cells were blocked and permeabilized with 3% donkey serum (Millipore) and 0.20% Triton-X100 (Sigma Aldrich) in TBS for 30 mins at room temperature. They were then stained with different combinations of the following primary antibodies: Goat α-GFP (1:1000, Rockland; Cat# 600-101-215), chicken α-GFP (Aves; Cat# GFP.1020), Mouse α-Lamin A (Millipore; Cat# MAB3540), Mouse α-mono and poly ubiquitin (Enzo Life Sciences; Cat# BML-PW8810-0500). The primary antibodies were incubated overnight at 4°C in the blocking buffer. The following day, the cells were washed 3 times with TBS. Then, the cells were incubated with the appropriate secondary antibodies for 2 hours at room temperature in the blocking buffer. Following this, the cells were washed 3 times with TBS and stained against DAPI. The coverslips were then mounted ImmuMount (Thermo Fisher Scientific) and imaged.

### Statistical tests

All statistical tests were performed using Prism. The graphs plotted show mean ± standard error of mean unless otherwise stated. The p values are denoted by the following: ns > 0.05, *p < 0.05, **p < 0.01, ***p < 0.001.

## Acknowledgments

This work was supported by the European Research Council (STEMBAR to S.J.), the Swiss National Science Foundation (BSCGI0_157859 and 310030_196869 to S.J.), and the Zurich Neuroscience Center. We thank Roche for providing us with hESC-derived NPCs. We also thank Dr. Magdalini Polymenidou and Dr. Marian Hruska-Plochan for graciously providing iPSC-derived NPCs.

## Author Contributions

M.K.B.I performed experiments, analyzed data, and co-wrote the manuscript. L.N.R performed experiments and edited the manuscript. S.J. developed the concept and co-wrote the manuscript.

## Supplementary figures

**Figure S1.**
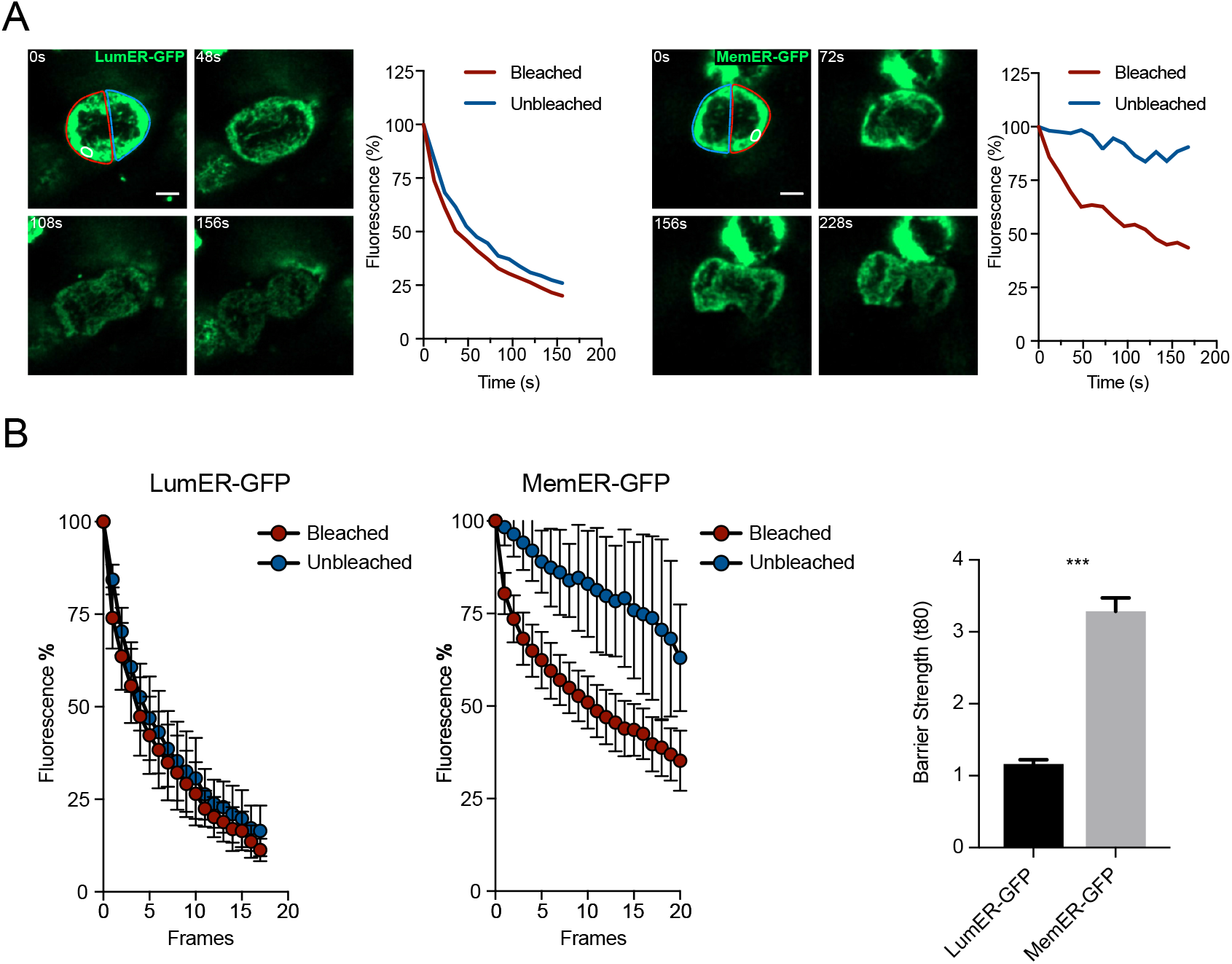
ER membrane diffusion barrier is formed in a second line of hNPCs. Related to Figure 1. (A) Single cells expressing either LumER-GFP (right) or MemER-GFP (left) undergoing FLIP assays are shown. The region bleached is highlighted (white). The Bleached and the Unbleached compartments of the cells are shown (red and blue respectively). Measured fluorescence intensities for the two compartments are plotted against time (right). (B) Average fluorescence intensities for the Bleached and the Unbleached regions are shown for LumER-GFP and MemER-GFP (left). Barrier strength indices of both MemER-GFP and LumER-GFP are also plotted (right). Scale bars represent 5μm (A). ***p < 0.001

**Figure S2.**
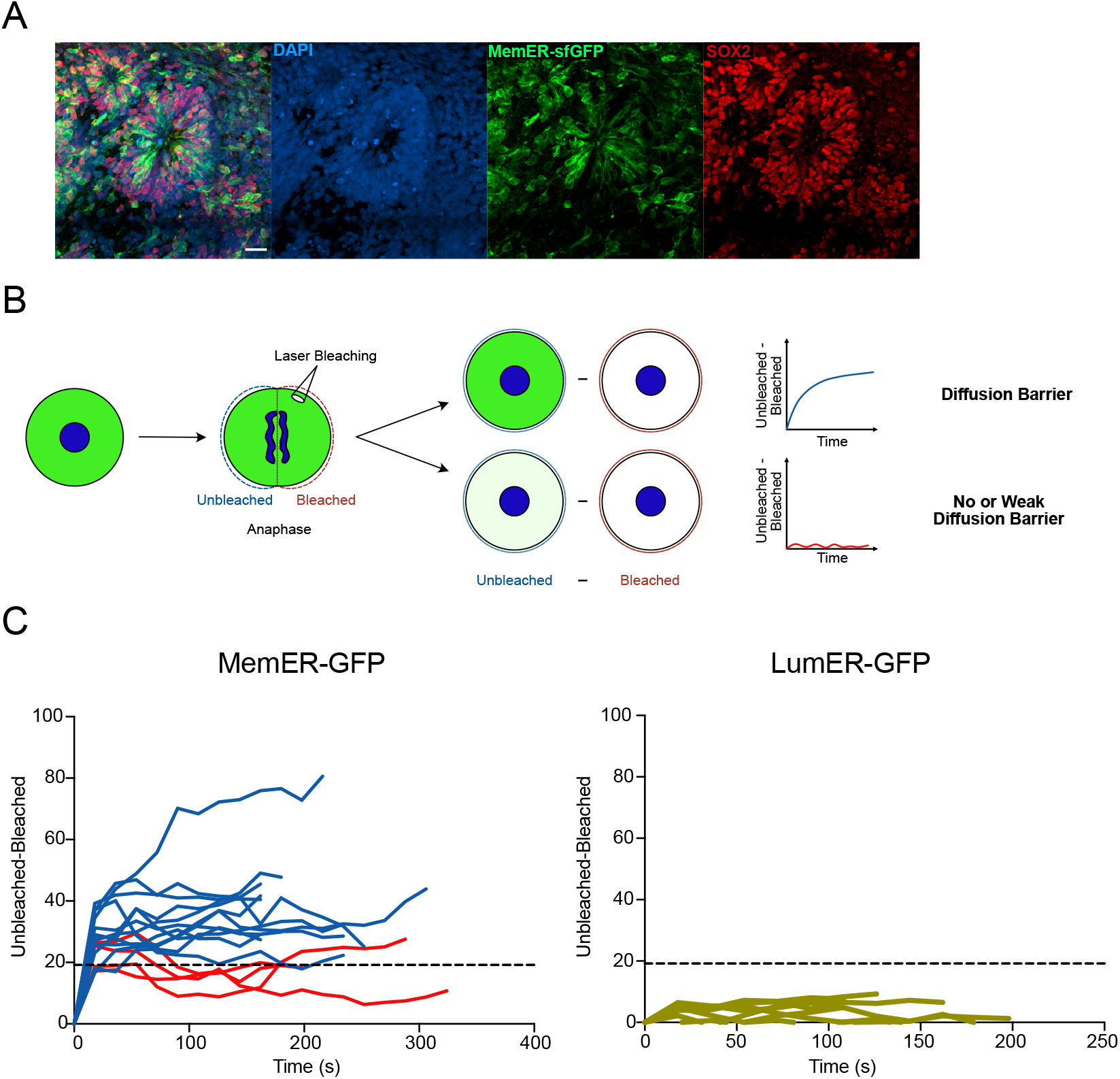
Neural progenitors in human forebrain organoids establish an ER membrane diffusion barrier. Related to Figure 4. (A) Human forebrain organoids stained against neural progenitor markers are shown. DAPI signal (blue), MemER-GFP (green) and SOX2 (red) is shown. (B) Outline for the difference analysis is shown. The cell undergoes FLIP as described before. Then, the bleached intensity is subtracted from the unbleached intensity at each timepoint. In the presence of a diffusion barrier, the difference between the unbleached and bleached would be high (blue) whereas in the absence of a barrier, the difference between the two will be low (red). (C) Difference analysis for each cell in the human forebrain organoids that had underwent FLIP assays is shown. For Lum-ER-GFP, all cells showed a weak or no diffusion barrier (red). For MemER-GFP, 4 cells showed a weak or no diffusion (red) barrier whereas 12 cells showed a diffusion barrier (blue). The dotted line above cells were classified as having a barrier is at y=19.18, which is the standard deviation of the LumER-GFP averaged over all the timepoints and multiplied by 6. Scale bars represent 20μm (A).

**Movie S1:** FLIP experiments on human NPCs expressing either LumER-GFP or MemER-GFP are shown. The human NPCs were derived from either human iPSCs or human ESCs, as indicated in the movie. Bleached area (white region) is shown exactly in the first frame and approximated in the subsequent frames. Scale bar represents 5μm.

**Movie S2:** FLIP experiments on human NPCs expressing MemER-GFP with either IRES-CFP (control) or Progerin-IRES-CFP (Progerin) are shown. The NPCs were derived from human ESCs. Bleached area (white region) is shown exactly in the first frame and approximated in the subsequent frames. Scale bar represents 5μm.

**Movie S3:** FLIP experiments on human forebrain organoids expressing LumER-GFP or MemER-GFP are shown. Bleached area (white region) is shown exactly in the first frame and approximated in the subsequent frames. Scale bar represents 5μm.

